# Individual-Based Integral Projection Models: The Role of Size-Structure on Extinction Risk and Establishment Success

**DOI:** 10.1101/029165

**Authors:** Sebastian J. Schreiber, Noam Ross

## Abstract

1. Matrix models or integral projection models (IPMs) are commonly used to study the dynamics of structured populations, where discrete or continuous traits influence survival, growth, or reproduction. When a population’s size is small, as is often the case for threatened species or potentially invasive species arriving in novel habitats, extinction risk may be substantial due to demographic stochasticity.
2. Branching processes, which are individual-based counterparts to matrix models and IPMs, allow one to quantify these risks of extinction. For discretely structured populations, the theory of multi-type branching processes provides analytic methods to compute how extinction risk changes over time and how it depends on the size and composition of the population. Building on prior work on continuous-state branching processes, we extend these analytic methods to individual-based models accounting for any mixture of discrete and continuous population structure.
3. The individual-based IPMs are defined by probabilistic update rules at the level of the individual which determine how each individual with a given trait value dies, changes trait value (e.g. grows in size), or produces individuals with the same or other trait values. Probabilities of extinction are shown to be analytically determined by probability generating functionals associated with the individual-based IPMs. In particular, we present analytical expressions for how extinction probabilities change over time and depend on the initial abundance and trait distribution of the population. We illustrate how to numerically implement these methods using data from the short-lived desert shrub species *Cryptantha flava*, and provide a more general discussion of how to implement these methods to other data sets including those involving fluctuating environmental conditions.
4. As most IPM studies have the necessary data to parameterize individual-based IPMs, these methods provide a computationally efficient means to explore how continuously structured populations differing in their evolutionary history and environmental context may differ in their vulnerability to extinction or ability to colonize new habitats.

## Introduction

Computations of extinction probabilities or likelihoods of establishment success lie on opposing sides of a theoretician’s coin and have been used to address theoretical and practical issues in conservation biology, restoration ecology, biological invasions and population genetics. Risks of extinction or establishment failure stem from populations consisting of a finite number of individuals, each of which faces a non-zero risk of mortality on any given day. These extinction risks are shaped, in part, by the size and composition of a population whose individuals may differ in age, size, geographical location, or other important characteristics influencing demography. When population structure is finite-dimensional (e.g. a finite number of age classes, stages, geographical locations), multi-type branching processes can model these extinction risks and, thereby, serve as the stochastic, finite-population counterpart of matrix models [Harris, 1963, Athreya and Ney, 2004, Caswell, 2001, Haccou et al., 2005]. These stochastic models have been used successfully to address a diversity of questions concerning fixation probabilities of beneficial alleles [Patwa and Wahl, 2008], evolutionary emergence of pathogens [Antia et al., 2003, Park et al., 2013], extinction risk of small populations [Boyce, 1992, Gosselin and Lebreton, 2000, Fujiwara and Caswell, 2001, Erickson et al., in press], and establishment success in heterogeneous environments [Haccou and Iwasa, 1996, Haccou and Vatunin, 2003, Schreiber and Lloyd-Smith, 2009].

To parameterize matrix models or multi-type branching processes, individuals must be discretely categorized into a finite number of types. However, when collecting demographic data, researchers commonly measure continuous traits (e.g. mass, length, geographical location) about individuals and use continuum-based statistics to approximate “fine-grained” discrete-traits. Integral projection models (IPMs) allow one to account for this continuous population structure [Easterling et al., 2000]. These IPMs can be viewed as infinite-dimensional matrix models and can be numerically approximated by finite-dimensional matrix models. Consequently, many of the standard demographic concepts and methods for matrix models (e.g. stable state distributions, reproductive values, life table response experiments, sensitivity analysis) exist for IPMs [Easterling et al., 2000, Ellner and Rees, 2006, 2007, Rees and Ellner, 2009, Coulson, 2012, Ellner and Schreiber, 2012, Metcalf et al., 2013, Rees et al., 2014, Merow et al., 2014].

Here, we describe individual-based counterparts of IPMs using continuous-state branching processes [Harris, 1963]. For these finite population, stochastic models, we present an analytical method for computing extinction probabilities. As these methods are easily implemented numerically, they circumvent the need to use individual-based simulations and allow one to efficiently study how extinction probabilities or establishment failure depend on continuous as well as discrete population structure. We illustrate the application of these methods with an individual-based IPM of the short-lived desert shrub *Cryptantha flava* from Utah, USA [Salguero-Gómez et al., 2012].

## The General Models and Methods

**The Individual Based IPM.** We consider an individual-based model where the set of all possible individual states (e.g. age, size, geographical location, etc.) lies in a compact metric space *X*. For a standard size-structured IPM, *X* = [*a,b*] corresponds to the range of sizes measured in the field where *a* is the minimal size and *b* is the maximal size. For models with a mixture of age and size structure, *X* could be given by {1,… *,T} ×* [a, b] where *T* corresponds to the maximal age of an individual.

Following Harris [1963], we consider finite populations in which the state of the population at any point in time is characterized by the different states (*x*_1_, *x*_2_, …, *x_k_*) of individuals within the population and the number of individuals in each state (*n*_1_*,n*_2_, …, *n_k_*). Specifically, if there *n*_1_ individuals in state *x*_l_, *n*_2_ individuals in state *x*_2_, …, *n_k_* individuals in state *x_k_*, then the state of the population is given by s = (*n*_1_, *n*_2_, …, *n_k_*; *x*_l_, *x*_2_,…, *x_k_*). The set of all possible population states is

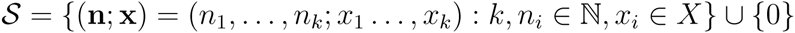

where ℕ = {1, 2, 3,… } denotes the natural numbers and 0 is the extinction state corresponding to no individuals in the population.

Let 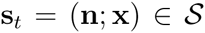 be the population state at time *t*. The dynamics of s*_t_* are determined by a set of probabilistic rules that determine the contribution of each individual in the population to the population in next time step *t* + 1. These “contributions” may correspond to an individual surviving and changing state (e.g. growing in size, getting older, dispersing to another geographical location), or an individual having offspring. Consistent with standard branching process theory, each individual updates independently of all other individuals in the population.

The update rule for an individual in state *x* is given by a probability measure m(*x*, *d*s) on the state space 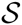. Specially, the probability an individual in state *x* contributes **s** individuals to the population in the next time step where **s** lies in a subset 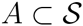 equals

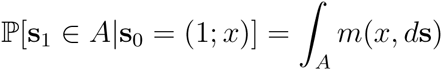

where the left hand side reads “the probability the population state lies in *A* at time 1 after initially having only one individual in state *x* at time 0.” If the population state is currently s*_t_* = (*n*_1_,…, *n_k_*; *x*_1_,…, *x_k_*), then the state **s***_t_*_+l_ is determined as follows:

1. for each of the *n*_1_ individuals in state *x*_1_, randomly and independently choose the number of replacement individuals from distribution *m*(*x_1_,d***s**),
2. repeat step (1) for the states *x*_2_, …, *x_k_*, and
3. determine the new population state s*_t_*_+1_ by identifying the states of all individuals and counting the total number of individuals in each of these states.

This iterative algorithm can be used to create individual based simulations of the individual based IPM. As with any branching process, stochastic realizations of this process, with probability one, either go to extinction in finite time, or the population abundance grows without bound. This latter event is typically interpreted as a population becoming established or persisting.

**Probability generating functionals and extinction probabilities**. We can characterize the probabilistic state of the system using probability generating functionals Ψ (pgfs). Unlike moment generating functionals as used by Harris [1963], the pgfs allow us to directly compute how extinction probabilities change in time as well as compute asymptotic extinction probabilities.

To define the pgf Ψ, we introduce the following notation: given a continuous function *h*: *X* → ℝ and 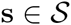 let

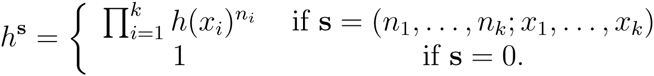

The utility of this definition stems from the observation that if *h*(*x*) corresponds to the probability that an individual of size *x* dies and has no offspring over the next year, then *h^s^* corresponds to the probability that a population in state **s** goes extinct in the next year. The probability generating functional is defined by

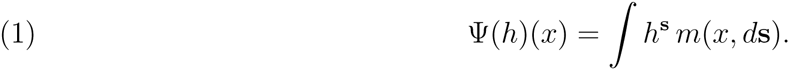

Ψ(*h*)(*x*) corresponds to the expected value of *h*^s^ due to the contributions of individuals in the next time step from an individual in state *x*. This expected value requires integrating over all possible populations states in the next time step.

The utility of Ψ for computing extinction probabilities follows from two facts. First, the definition of Ψ implies that if h_0_ is the zero function (i.e. *h*_0_(*x*) = 0 for all *x*), then

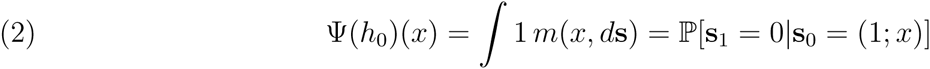

is the probability the population goes extinct in one time step given that initially it consisted of one individual in state *x*. For the second fact, define

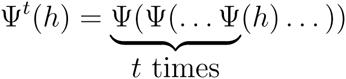

to be the *t*-fold composition of Ψ with itself. We claim that

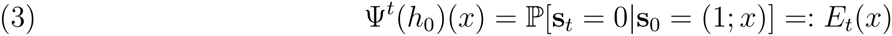

is the probability of extinction by time t given that the population initially consisted of one individual in state *x*. To verify this claim, we argue by induction. Equation 2 implies that equation (3) holds for *t* = 1. Now suppose that equation (3) holds at time *t*; we will show it holds at time *t* + 1. On the event that s_1_ = (*n*_1_, …, *n_k_*; *x*_1_, …, *x_k_*) is the population state at time 1, extinction occurs by time *t* + 1 only if each of the lineages of the n_1_ + ··· + n*_k_* individuals go extinct in the next *t* time steps. As the fates of these lineages are independent of one another, it follows that

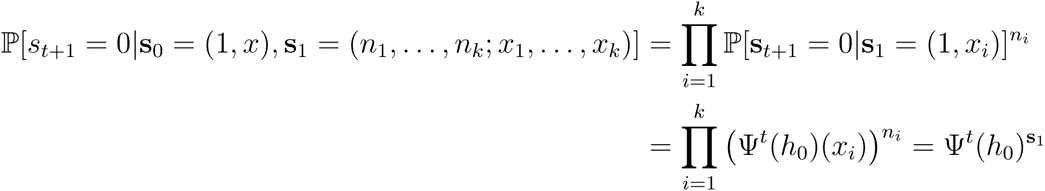

where the second equality follows from our inductive hypothesis. By the law of total probability

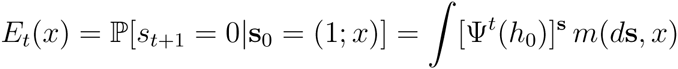

which, by definition, equals Ψ*^t^*^+l^(*h*_0_)(*x*) as claimed.

Equation (3) can be used to compute extinction probabilities iteratively. Furthermore, as individuals update independent of one another, the probability of the population going extinct by time *t* for any initial condition s = (*n*_1_,…, *n_k_*; *x*_1_,…, *x_k_*) equals

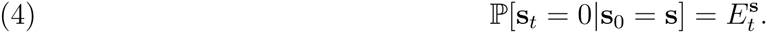

These analytic expressions allow us to efficiently compute extinction probabilities by constructing a numerical approximation of the pgf Ψ and iterating it with an initial condition of a zero vector which corresponds to the numerical approximation of the zero function.

As *E*_0_(*x*) ≤ *E*_1_(*x*), ≤ *E*_2_(*x*)… for any *x* ∈ *X* and *E_t_*(*x*) ≤ 1 for all *t*, there is a well defined limit corresponding to the probability of eventual extinction:

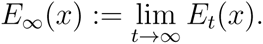

Using moment generating functionals with suitable technical hypotheses, Harris [1963] showed that *E_∞_*(*x*) < 1 for all *x* if the dominant eigenvalue of the mean-field IPM is greater than one, and *E_∞_*(*x*) = 1 for all *x* otherwise.

### An Illustration with a short-lived perennial

We illustrate these general methods using an individual-based IPM for the yellow-flowered perennial plant, Plateau Yellow Miner’s Candle (*Cryptantha flava*), of the borage family *(Boraginaceae*). For populations growing along the Colorado Plateau, USA, Salguero-Gómez et al. [2012] developed an IPM using data collected from 2004 to 2010. Here, we use a subset of this data available in an R package, IPMpack [Metcalf et al., 2014]. All code to for this example is archived at Zenodo [Schreiber and Ross, 2015].

In the model, the state *x* of an individual is the size, which equals the total number of vegetative and flowering rosettes. If *N_t_*(*x*) denotes the density of individuals of size *x* in year *t*, then Salguero-Gómez et al. [2012] used an IPM of the form

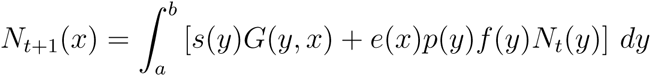

where *s*(*y*) is the probability of surviving to the next year for individuals of size *y*, *G*(*y, x*) *dy* is the infinitesimal probability that a surviving individual of size *y* is size *x* in the next year, *p*(*y*) is the probability that an individual of size y flowers, *f*(*y*) is the mean number of offspring produced by an individual of size *y*, and *e*(*x*) is the infinitesimal probability that an offspring is of size *x* at the time of the annual census.

Following Salguero-Gómez et al. [2012], we use generalized linear models (GLMs) for most of the functional forms of the demographic kernels. Computations were performed using the base GLM function in R [R Core Team, 2015]. We used logistic regression (i.e. a GLM with the binomial family) for determining *s*(*y*) and *p*(*y*), and a GLM with the Poisson family for modelling *f*(*y*). For the growth kernel, linear regression determined the expected size of an individual in the next year and the actual size was assumed to be normally distributed about this mean. The variance of this normal distribution, for simplicity, was assumed to be independent of the current size of an individual. We modeled *e*(*y*) with a gamma distribution fit to the the empirical distribution of germinants. Figure 1 shows the data and fits for *s*(*y*), *G*(*y,x*), *p*(*x*) and *f*(*x*).

**Figure 1.**
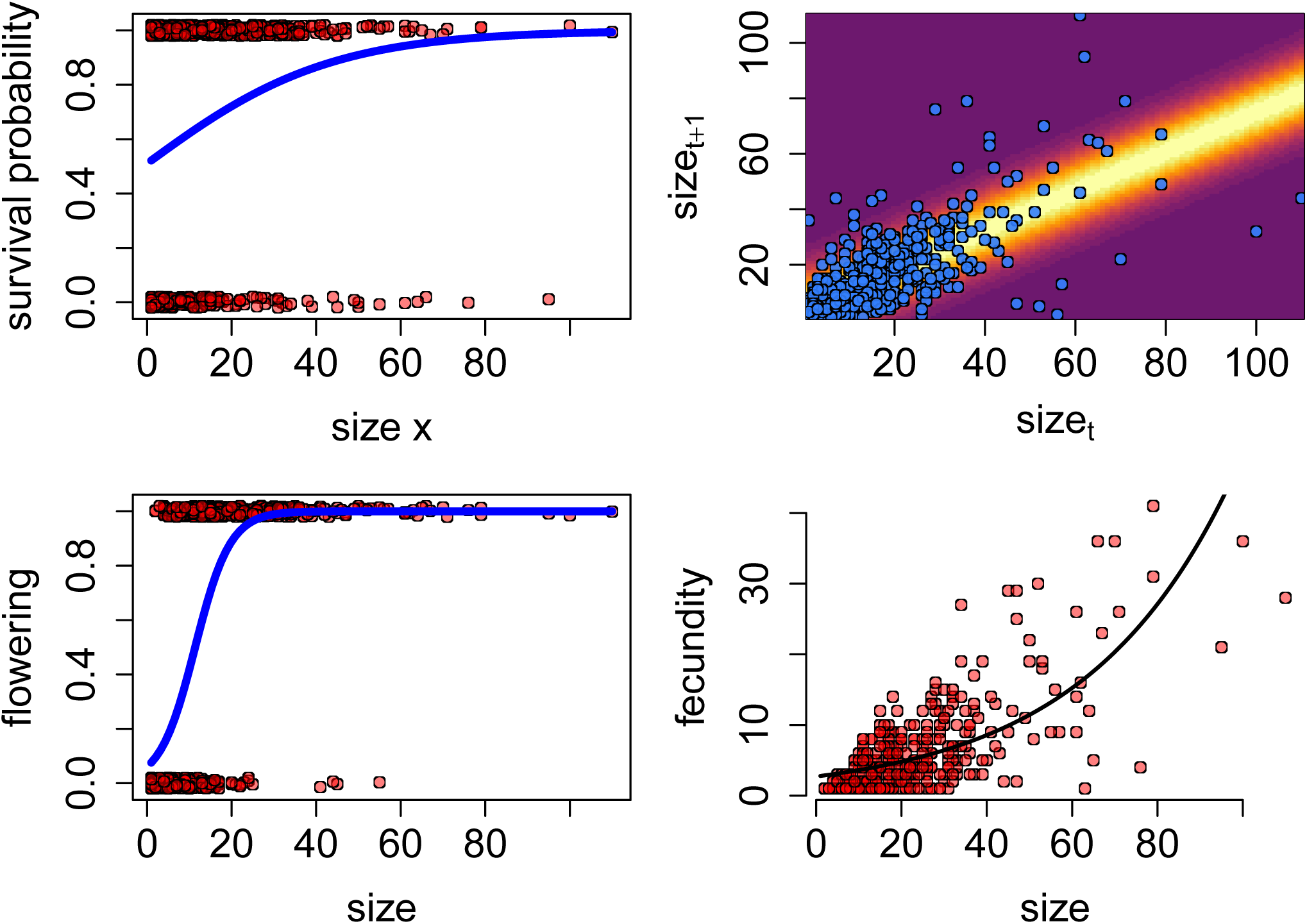
The demographic kernels for Plateau Yellow Miner’s Candle with the corresponding data from Salguero et al. 2012.

The kernels *s*(*x*), *G*(*y,x*), *p*(*x*) and *e*(*x*) provide us with all the information that the individual-based IPM requires for probabilistic updating individuals for survival, growth, flowering, reproduction, and size of germinating individuals at first census. The fecundity kernel *f*(*y*), however, only specifies the mean number of offspring produced, but for an individual-based IPM, we need the distribution of the number of offspring produced by an individual. Fortunately, this information is built into the structure of the GLMs due to the assumptions in our model choice. As the fecundity data was modeled using a Poisson family for the GLM, the mean number *f*(*x*) of offspring also specifies the distribution. More generally, one might use multi-parameter distributions such as a zero-inflated Poisson or a negative binomial, in which case parameters in addition to the mean are needed to specify the distribution of offspring number.

#### Deriving the pgf Ψ

To define Ψ, we observe that the contributions of an individual of size *x* to the population in the next time step involves the sum of two independent random variables: the contribution due to survival and growth and the contribution due to reproduction. We will identify two pgfs, Ψ*_g_* and Ψ*_f_*, for each of these processes separately. Then, we make use of a fundamental property of pgfs

##### Fundamental Property 1

The pgf for a sum of independent random variables is the product of the pgfs of these random variables.

to get that

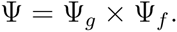

To write down each of these pgfs, we make use of another fundamental property of pgfs:

###### Fundamental Property 2

The pgf for a sum of *N* independent, identically distributed random variables *X_i_* is the composition of the pgf for *N* with the pgf for the *X_i_*.

For survival and growth, Ψ*_g_* (*h*)(*x*) corresponds to integrating *h*^s^ over all possible contributions **s** from an individual of size *x* surviving and growing. These contributions are of two types: **s** = 0 when the individual dies, and **s** = (1; *y*) when the individual survives and grows to size *y*. The first event occurs with probability 1 − *s*(*x*) and the infinitesimal probability of the second event is *s*(*x*)*G*(*y, x*)*dy*. As *h*^s^ = 1 when **s** = 0 and *h*^s^ = *h*(*y*) when **s** = (1; *y*), integrating over all possible contributions due to survival and growth yields

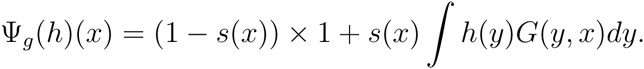

For fecundity, Ψ*_f_* (*h*)(*x*) is given by integrating *h*^s^ over all possible states **s** corresponding to the offspring produced by an individual of size *x*. To write this down, we begin by conditioning on the event that an individual of size *x* flowers. On this event, the individual produces a Poisson number *N* of offspring with mean *f*(*x*). The pgf for *N* is given by ø(*x, ξ*) = exp(− *f*(*x*)(ξ − 1)) where ξ is a dummy variable. The size of each of these offspring is drawn interdependently from the common offspring distribution *e*(*y)dy*. Hence, the contribution of a flowering individual of size *x* is the sum of *N* independent random variables with distribution *e*(*y)dy*. By **Fundamental Property 2** of pgfs, we need to take the composition of the pgf *φ* for *N* with the pgf for a single offspring, namely

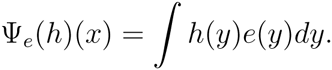

Thus, we get the pgf associated with a flowering individual is

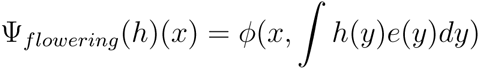

To get the pgf for flowering and non-flowering contributions to fecundity, we observe that fecundity contributions of an individual of size *x* is given by the sum of a Bernoulli number of flowering individuals where the probability of success is *p*(*x*). By the **Fundamental Property 2**, we need to compose the pgf of a Bernoulli, which is θ(ξ) = 1 − *p*(*x*) + *p*(*x*)ξ where ξ is the dummy variable, with the pgf Ψ*_flowering_* of a flowering individual. This composition yields

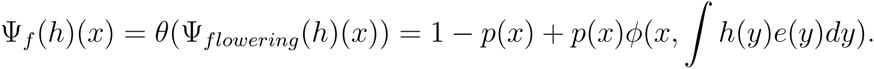

Muliplying Ψ*_g_* and Ψ*_f_*, we get the desired pgf Ψ:

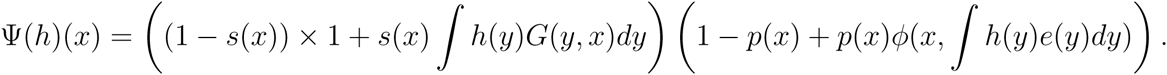

#### Numerically Implementing and Using the pgf Ψ

To approximate Ψ. numerically, we discretize a larger size interval [*a*, 2*b*] using *n* = 500 equal sized intervals of width Δ*x* = (2*b* − *a*)/*n*. We used the larger size interval of [*a*, 2*b*] to minimize the effects of eviction (see below). We created a vector 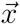 corresponding to the midpoints of these intervals. Using this vector we discretized the survival function as a vector 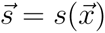 the growth kernel as a matrix using the outer product *G* of the growth kernel *g* with 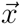 the probability of flowering function as a vector 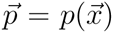 the fecundity function as a vector 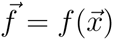 and the offspring size distribution as a vector 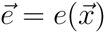

For the previously described methods to work, it is critical that column sums for *G* and the sum of 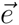 equal one. For most IPMs, this will not occur automatically due to individuals being evicted from the size interval [*a*, 2*b*]. There are a variety of ways to handle this issue [Williams et al., 2012]. As the offspring size vector nearly summed to one, we simply re-normalized it so that it summed to one. For the growth matrix *G*, we treated eviction as mortality. To do this, we took one minus the column sums of *G*, subtracted these sums from the survival vector, and then normalized the column sums of *G* so that they added to one. When taking the product of survival and growth, the resulting mean-field IPM is unaffected by these changes.

Using these discretized demographic components and the pgf *φ* for fecundity, we get the discretized pgf Ψ, given by

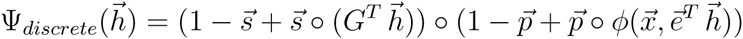

where 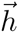 corresponds to a discretized function i.e. a vector of length *n, ^T^* denotes the transpose of a matrix or vector, and ○ denotes element by element multiplication.

Iterating Ψ yields how extinction probabilities *E_t_*(*x*) vary with time for a population initiated with a single individual (Figure 2). Intuitively, this figure illustrates that the probability of extinction decreases with the size of founding individual, and that extinction probabilities increase over time. Furthermore, *E_t_*(*x*) as *t* increases are approaching limiting extinction probabilities *E_∞_*(*x*) which always equals one (Figure 2A). This stems from the fact that the dominant eigenvalue of the mean-field IPM is less than one, consistent with the results of Salguero-Gómez et al. [2012]. By increasing seed survivorship by a factor of three, the dominant eigenvalue of the mean-field IPM becomes greater than one and the asymptotic extinction probabilities become less than one (Figure 2B). We approximated these asymptotic extinction probabilities by iterating Ψ until the difference between *E_t_* and *E_t_*_+1_ were below a specified error tolerance.

**Figure 2.**
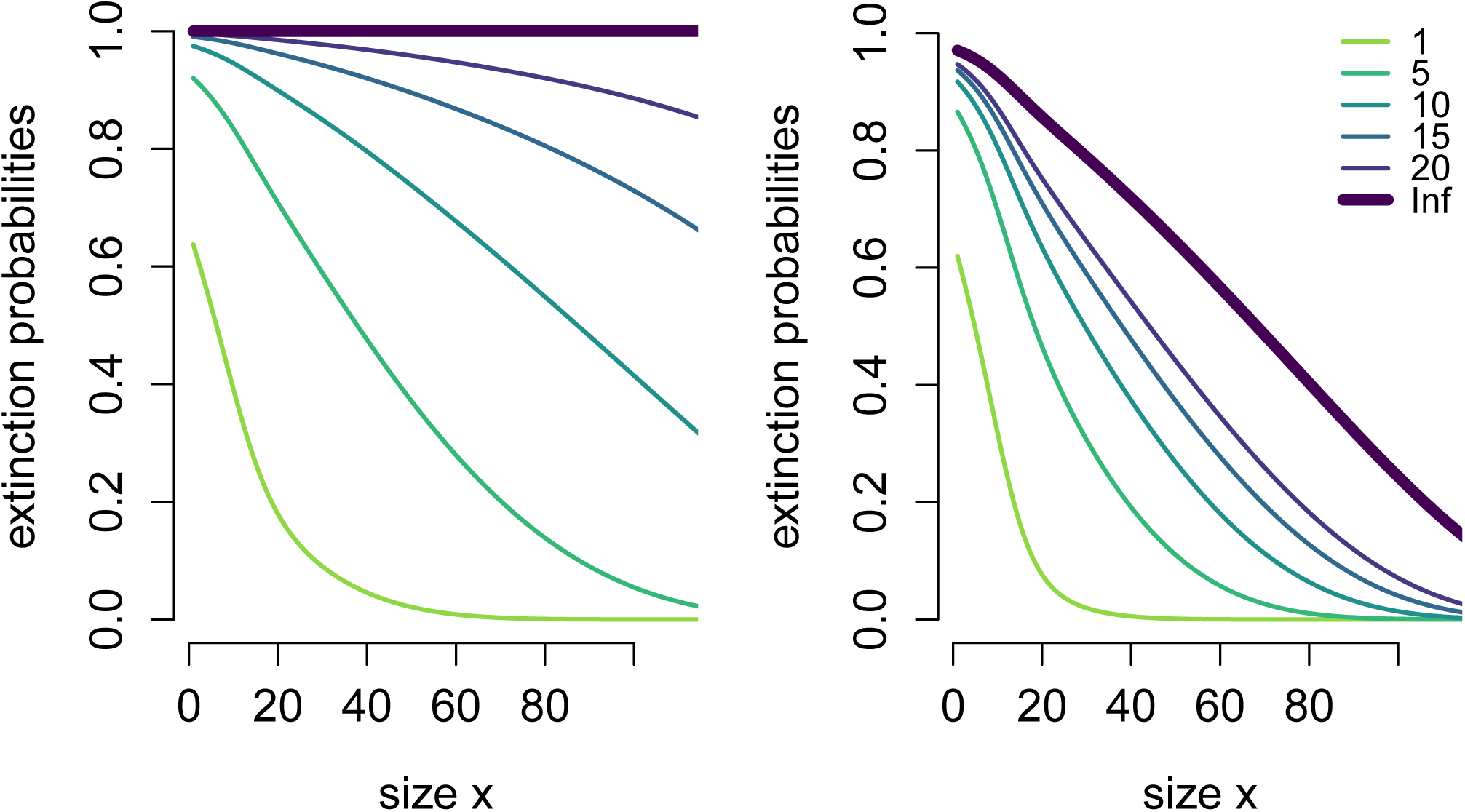
Extinction probabilities for a population initially consisting of a single individual of size *x*. Different curves correspond to extinction occuring in 1, 5, 10, 15 or 20 years. Asymptotic extinction probabilities are shown by the thicker curve. In A, the extinction curves for the baseline individual-based IPM. In B, extinction curves for the case when seed surival is increased by a factor of three.

To scale things up to an entire population initially in state s_0_, recall that the extinction probability at time *t* is 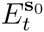 Figure 3 illustrates how the extinction probabilities over time vary for a population with initially 100 individuals of the smallest size *x* =1 (i.e. s_0_ = (1; 100)) versus a population with 5 or 8 individuals of the largest size *x* = 60 typically observed in the field (i.e., s_0_ = (60; 5) or (60; 8). Figure 3 suggests that, from the extinction risk perspective, about 6 or 7 larger individuals are equivalent to 100 of the smallest individuals. These types of comparisons may be particularly useful when trying to assess whether planting small or larger individuals are more effective for establishment success.

**Figure 3.**
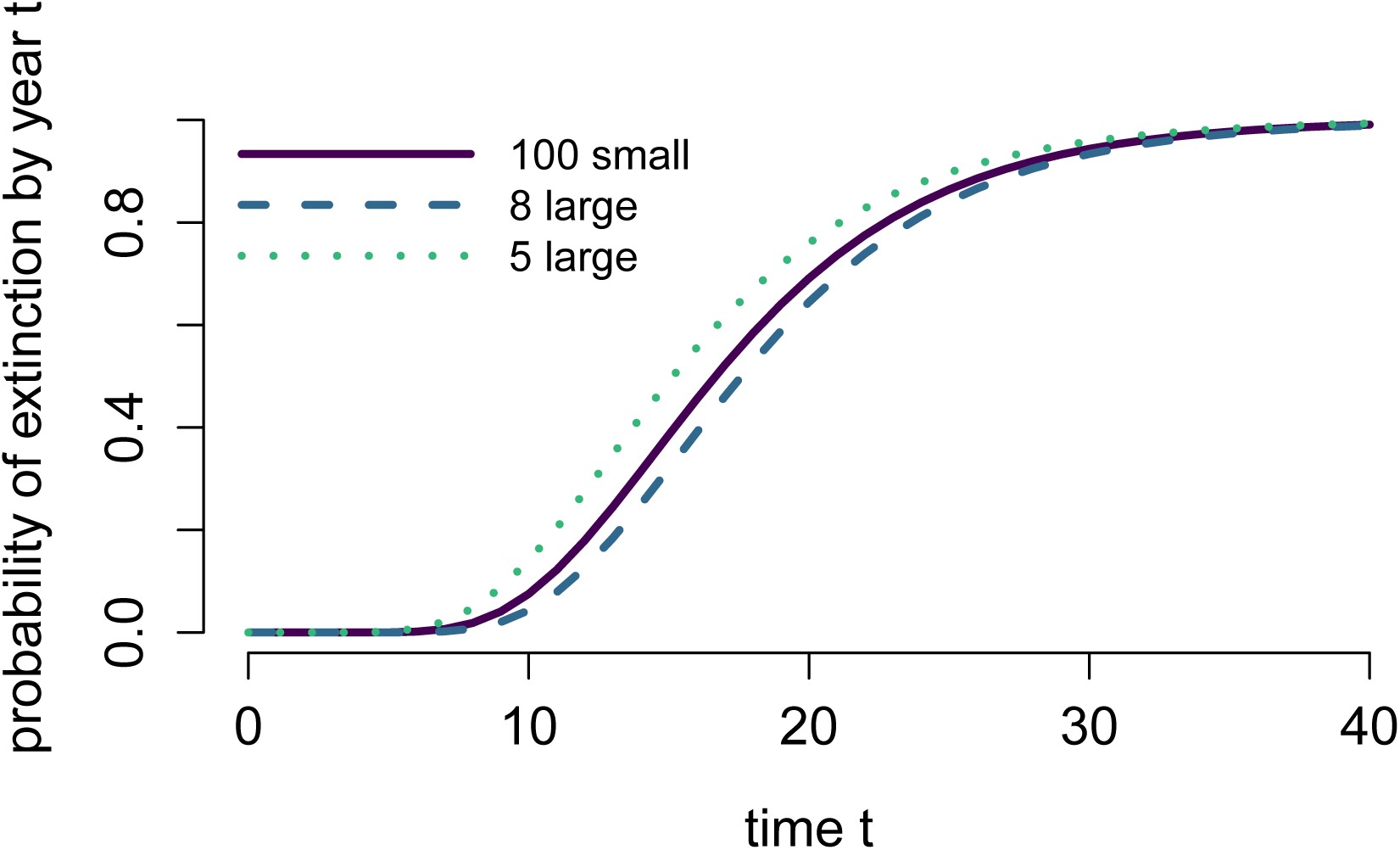
Extinction probabilities as a function of time for populations with 100 individuals of size *x* =1, and populations with 5 or 8 individuals of size *x* = 60.

We also used the data to see how extinction risk over different time frames depends on the size and composition of the population. Specifically, for different founding population abundances *N*, we randomly sampled *N* individuals from the data and computed extinction risk of this sampled population over 5 and 10 year periods (Figure 4). For each founding population abundance *N*, we considered 500 random samples of size *N.* Figure 4A illustrates that, on average, log extinction risk decreases with the founding population size *N* and is greater for the 10 year period than the 5 year period. For smaller founding popualation sizes *N*, there is substantial overlap in the distributions of extinction times for the 5 and 10 year time frames. This overlap stems from founding populations of mostly large individuals being more likely to persist at least 10 years than founding populations of mostly small individuals persisting at least 5 years. Consistent with this explanation, Figure 4B illustrates that mean size of an individual within a founding population has a strong negative correlation with (log) extinction probability, and the slope of this correlation is steeper over shorter time frames than longer time frames.

**Figure 4.**
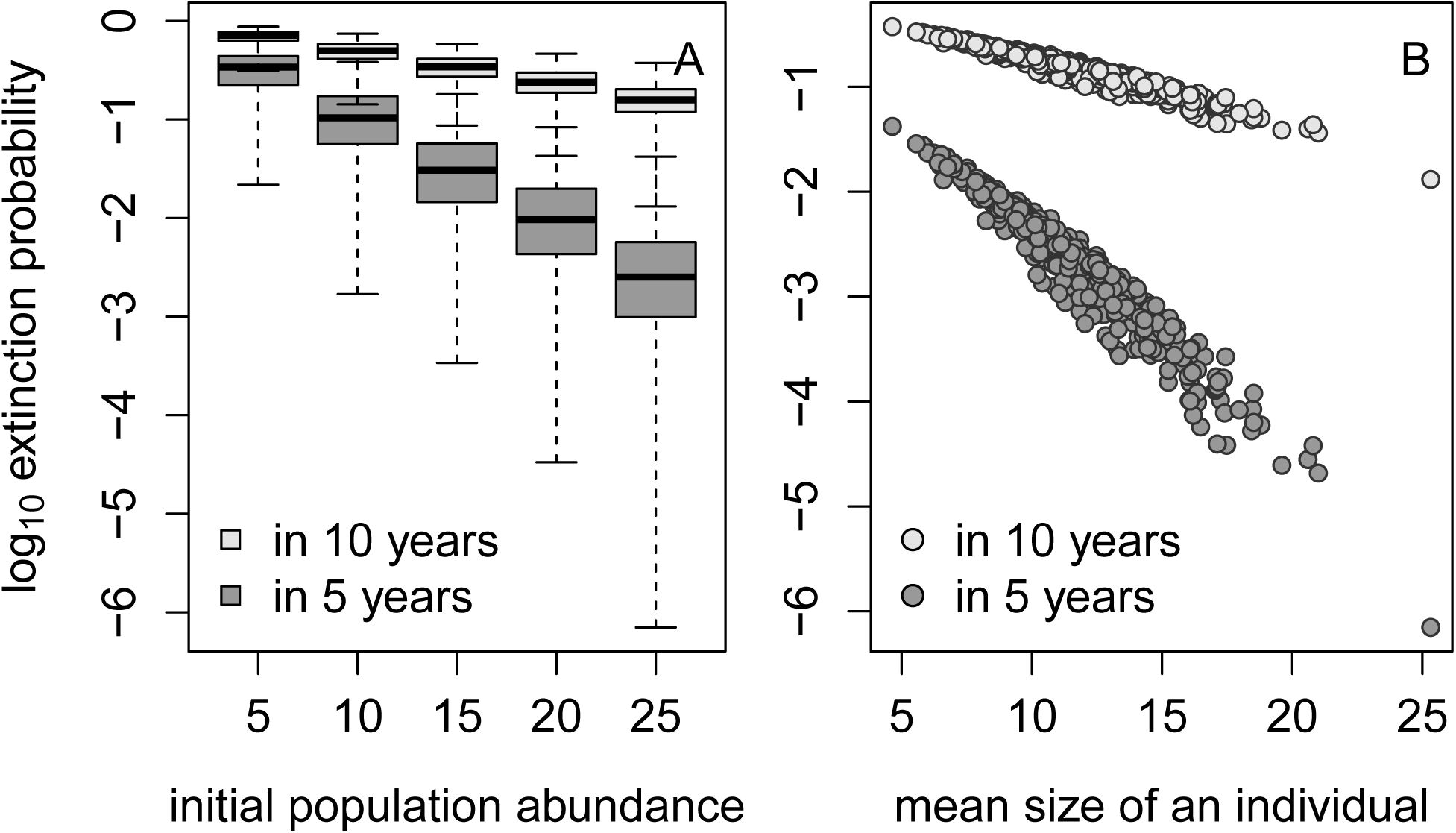
Extinction probabilities for founding populations of different abundance. For each founding population abundance *N*, 500 samples consisting of *N* randomly choosen individuals from the data set were used to create a founding population of *N* individuals. Extinction probablities by year 5 and year 10 were calculated for each of these sample populations. In A, extinction probability is plotted as a box plot for different *N* values. In B, extinction probability is plotted against the mean size of an individual for a population abundance *N* = 25.

### Recommendations, Extensions, and Future Challenges

To implement the methods presented here, there are two main steps. First, one needs to identify the main demographic processes of the population, the order in which these processes occur relative to the censuses used for data collection, and develop the statistical models for the each of the demographic processes. Rees et al. [2014] and Merow et al. [2014] provide excellent reviews on the philosophical and methodological issues associated with this step. Typically, whenever a study has sufficient data for constructing the mean-field IPM, there is no need to collect any additional data to build the individual-based IPM. Unlike the mean-field IPMs, though, the individual-based IPMs make use of the complete distributional information associated with fecundity. As with other areas of stochastic demography, the shapes, not just the means, of these distributions may have significant effects on the likelihood of extinction or establishment success [Lloyd-Smith et al., 2005]. Hence, it is best to examine several options (e.g. Poisson versus negative Binomial versus zero-inflated distributions) to identify which distribution does a better job of describing the fecundity data. As in all areas of modeling, if there is significant uncertainty about the “best” choice of the model, one should perform the analyses with each of the alternative fecundity distributions to identify the sensitivity of predictions of extinction risk to these alternatives.

The second step, the focus of this paper, involves constructing the probability generating functional Ψ. For the uninitiated, this step may be intimidating. However, there are a three basic principles that simplify this construction. First, while this pgf Ψ takes functions to functions, one should focus on writing down Ψ(*h*)(*x*), which involves understanding the contributions of a single individual of size *x* to the next census. Second, one can often can break up these contributions into a sum of independent contributions, find the pgfs associated with these simpler contributions, and then use the two fundamental properties of pgfs to “stitch” together Ψ. Third, the distributions used to describe the number of offspring produced by an individual typically involve random variables (e.g. Poisson, negative binomial) for which the associated pgfs are well-known. Finally, whenever in doubt, find a collaborator that you trust to help put the pieces together correctly.

For the individual-based models considered here, we assumed the environment remains constant over time. However, IPMs have and continue to be used to study the effects of fluctuating environmental conditions on population demography and life history evolution [Childs et al., 2004, Dahlgren and Ehrlen, 2011, Rees and Ellner, 2009]. The methods described here easily extended to fluctuating environments. Specifically, if Ψ*_t_* is the pgf of the individual-based IPM associated with year t, then the probability of extinction for a population initially in state **s** is

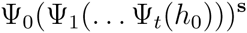

where *h*_0_, as before, is the zero function. Note that the composition here is in the reverse order of what one does when iterating the mean-field IPM model forward in time. While we know of no formal proof, in the case of a stationary environment with suitable technical assumptions, we conjecture the following limit theorem holds: the asymptotic extinction probabilities are strictly less than one if and only if the stochastic growth rate (aka dominant Lyapunov exponent) of the mean-field IPM is positive. For multi-type branching processes, this result was proven by Tanny [1981].

Despite this relatively straightforward extension to temporally variable environments, many challenges remain. From the computational perspective, finding the efficient methods to deal with multi-dimensional states variables (e.g. size and location, or multi-dimensional traits) continues to be a challenge, as it is for mean-field IPMs. While the analytical methods presented here cover multi-dimensional state variables, their numerical implementation involves approximating mutli-dimensional integrals which can be computationally expensive. From the analytical perspective, accounting for temporal correlations in individual growth or reproductive rates (e.g. individuals that grew larger than expected in one year being more likely to grow larger than expected in the next year) or correlations among individuals are particularly important challenges as strong correlations likely have large effects on extinction risk. Finally, and perhaps most importantly, the manners in which size-structured demography may shape extinction risk for real-world population remains to be understood. One might hope that by applying these methods to the many data sets for which IPMs have been developed, as well as future data sets, might provide a computationally efficient means to explore how size-structure for populations differing in their evolutionary history and environmental context influences their vulnerability to extinction.

## Acknowledgements.

We thank William Cuello, Eric Eager, Richard Erickson, Vadim Karatayev, and Jacob Moore for providing comments on an earlier draft of this manuscript.

